# A mesocorticolimbic signature of pleasure in the human brain

**DOI:** 10.1101/2022.10.31.514244

**Authors:** Philip A. Kragel, Michael T. Treadway, Roee Admon, Diego A. Pizzagalli, Emma C. Hahn

**Author notes:** Please address correspondence to: Philip A. Kragel, Department of Psychology, PAIS 391 Emory University, Atlanta, GA 30032.

## Abstract

Pleasure is a fundamental driver of human behavior, yet its neural basis remains largely unknown. Rodent studies highlight opioidergic neural circuits connecting the nucleus accumbens, ventral pallidum, insula, and orbitofrontal cortex as critical for the initiation and regulation of pleasure, and human neuroimaging studies exhibit some translational parity. However, whether activation observed across these regions reflects a common, generalizable code for pleasure driven by opioidergic mechanisms remains unclear. Here we use pattern recognition techniques to develop a human functional magnetic resonance imaging signature of mesocorticolimbic activity unique to states of pleasure. In independent validation tests, we find this signature has high sensitivity to pleasant tastes and positive affect evoked by humor. The signature is spatially coextensive with mu-opioid receptor gene expression, and its response is attenuated by the opioid antagonist naloxone. These findings provide evidence of a basis for pleasure derived from primary and secondary rewards in humans that is distributed across brain systems, and suggest that similar mechanisms underlie hedonic impact across mammalian species.

## Main

Pleasure is central to human experience, and has served as a cornerstone for philosophical^1,2^, socio-economic^3,4^, and psychological^5,6^ frameworks for understanding human behavior for thousands of years^7^. Despite its centrality for daily life and philosophical systems alike, the neuroscientific understanding of pleasure in the human brain remains in its infancy^8-11^. This stands in stark contrast to the study of human ‘reward’, which has largely focused on identifying neural systems that mediate behavioral responses to reinforcing stimuli^12,13^. Such paradigms have yielded crucial insights into the circuitry underlying conditioning, learning, and decision making, yet it is widely accepted that such behavioral manifestations of preference do not necessarily reflect the direct experience of pleasure^14^, and that pleasure is not necessary for reinforcement^6,13^. As such, the neural basis for subjective pleasure remains elusive.

Indeed, much of the known functional neuroanatomy of pleasure in mammals^10,11,15^ has been derived from studies in rodents, which have established a critical role for mu-opioid signaling within a network of regions including the nucleus accumbens shell, ventral pallidum, orbitofrontal cortex, and insula^11,16,17^. Critically, microinjections of mu-opioid agonists into specific zones in these areas, referred to as hedonic “hotspots”, enhance putatively pleasure-related behaviors involving relaxed facial expressions and rhythmic tongue and mouth movements^11,16-22^. These same injections also suppress pleasure-related behaviors in neighboring hedonic “coldspots”^18,22,23^. Importantly, the role of the mu-opioid system has been proposed as selective to pleasure-related behaviors and neurobiologically dissociable from putatively dopaminergic aspects of behavioral reinforcement, such as conditioning, craving, and invigoration^13,23^.

Attempts to translate this preclinical literature to humans have been mixed. On the one hand, human neuroimaging studies have shown that the same set of regions are commonly activated by diverse rewards^24-29^, with positron emission tomography further highlighting the involvement of endogenous opioids^30^. However, the functional homology of this network in humans and circuits identified in rodents is contested—particularly in prefrontal cortex and insula^31-33^. Moreover, studies of opioid antagonism on pleasure responses have revealed inconsistent effects, with some evidence suggesting that opioid antagonism may exhibit more effects on motivation than pleasure in humans^34,35^. Finally, the nature of pleasurable stimuli accessible for study in animal models does not extend to many important modalities of human pleasure, such as music^36^ and humor^37^. Consequently, the generalizability of rodent models for understanding the full complement of human pleasures is uncertain.

Uncertainty about hedonic brain systems in humans is also due to the size and spatial configuration of affective circuitry in subcortical structures, as well as the limits of conventional imaging approaches. In rodents, hedonic hotspots in the nucleus accumbens form a spatial gradient in which anterior areas are involved in appetitive and posterior areas in aversive behaviors^38^. A mirrored gradient is present in the ventral pallidum, in which activation of caudal areas enhances appetitive behaviors and rostral areas enhance avoidant behaviors^20^. These hotspots comprise only a small portion (∼10%) of the subcortical structures in which they are situated, which contain functionally and neurochemically heterogeneous neural populations^39-48^. Accordingly, when assessed with conventional fMRI, signals from different neural populations are blurred together, obscuring which affective variables (e.g., autonomic arousal and reward value) are encoded in each region^49-51^. This has limited efforts to characterize hedonic brain systems in humans, making it difficult to isolate neural substrates involved in different components of reward.

Due to these limitations, a growing number of researchers have turned to multivariate approaches to evaluate how affective variables are represented in human brain activity^52-55^. Unlike standard univariate analysis, multivariate methods are capable of estimating a spatial profile of activity within and across regions that most accurately characterizes a variable of interest^56,57^, even in cases where multiple neural populations overlap in a single region^58^. Indeed, pattern-based methods have revealed responses in orbitofrontal cortex and adjacent ventromedial prefrontal cortex that discriminate states of pleasure from displeasure^55^. Thus far, however, there is surprisingly limited evidence that neural populations in subcortical structures represent diverse pleasures using a common code, or that they form part of a distributed network mediated by opioidergic mechanisms positioned to influence learning and decision making as predicted by contemporary accounts of reward learning^13^. Moreover, because brain areas consistently engaged by rewards also respond to a wide array of motivationally salient, aversive, and painful stimuli^29,59^, it is unclear whether these regions contain circuitry that regulates pleasure across contexts as opposed to other nonspecific factors such as motivational salience or arousal^23,59,60^.

Understanding hedonic systems in humans has implications for translational research since altered regional activity in the ventral striatum, basal forebrain, and amygdala have emerged as candidate biomarkers for many psychiatric disorders^61,62^ and are common targets for clinical interventions^63-65^. Although specific regions of interest are often well motivated by preclinical research, recent work developing brain-based biomarkers of affective processes^66-69^ has shown that univariate measures often produce smaller effects^70^, and are less accurate and reliable than multivariate predictive models^71-73^.

Here we aim to more precisely model human brain responses to diverse pleasures, with a specific focus on regions known to contain hotspots in rodents. We combined a mega-analytic approach and pattern recognition techniques to characterize brain responses across 28 fMRI studies (total *n* = 494, see Methods). We used patterns of brain activity from 10 studies that manipulated pleasure using music, images of appetizing food, erotic images, cues of monetary rewards, and socially relevant stimuli (2 studies of each type, total *n* = 224) to develop an fMRI-based model, or brain signature, that predicts the hedonic state of an individual. Brain responses during manipulations of positive affect were differentiated from those acquired during affectively salient, but not pleasurable experiences (18 studies, n = 270)^74^.

Modeling brain activity across diverse experimental manipulations enabled us to tease apart signals that are consistent across instances of pleasure from those that are bound to a single stimulus or sensory modality^75^. By focusing on patterns of brain activity, we move beyond individual brain regions to characterize fine-grained topographies in distributed mesocorticolimbic circuitry that are defining features of hedonic systems. Training the model on data from 28 independent studies facilitated prospective tests of two key predictions from theories of hedonic function: 1) that a signature for pleasure should track the hedonic component of reward, as opposed to motivational or learning components, and further 2) that the response of such a signature should be mediated by opioid receptors. We verified that the signature was sensitive and specific to pleasure in four fMRI studies (total *n* = 89), two that evoked states of pleasure using primary reinforcers and two that manipulated prospects for subsequent monetary reward but had relatively less hedonic impact. Further, we evaluated opioid involvement in the signature response during a pharmacological challenge using the antagonist naloxone (*n* = 19). These validation tests provide a strong assessment of the hypothesis that human pleasure is mediated by a distributed opioidergic network.

### Towards a brain signature for pleasure

We used a latent variable multivariate regression technique, partial least squares regression^76^, to predict states of pleasure from patterns of fMRI activity. This approach estimates the spatial layout and activation of multiple latent sources that explain both observed fMRI activity and affective variables of interest^74^. The model included signals from brain regions known to contain hedonic hotspots and interconnected areas involved in emotion, motivation, and reward processing^77^ (see Methods for details). We identified a pattern across these areas that predicts pleasure independent of its sensory origin (whether music, images of food, erotic images, monetary reward, or social stimuli) while accounting for spatially overlapping signals that do not generalize across instances of pleasure or signals that are shared with other affective states.

The signature contained positive coefficients, which estimate the spatial layout of population activity associated with pleasure, in multiple regions including the anterior ventral insula, anterior agranular insula, midcingulate cortex, dorsomedial prefrontal cortex, basolateral amygdala, extended amygdala, globus pallidus, ventral striatum, and substantia nigra (voxel-wise FDR *q* < .05, see Fig. 1a and Supplementary Tables 1-2). Negative coefficients were present in posterior insula, dorsal mid-insula, midcingulate cortex, and supplementary motor area. In line with evidence of interdigitated population activity during appetitive and aversive behaviors^78-80^, most regions contained both positive and negative coefficients (Fig. 1b). Multiple regions differed in their balance of positive and negative coefficients. The habenula, anterior ventral insula, basolateral amygdala, internal globus pallidus, and midbrain regions contained more positive coefficients whereas middle and posterior insula contained primarily negative coefficients (*q*_FDR_ < .05, Supplementary Table 2).

**Figure 1.**
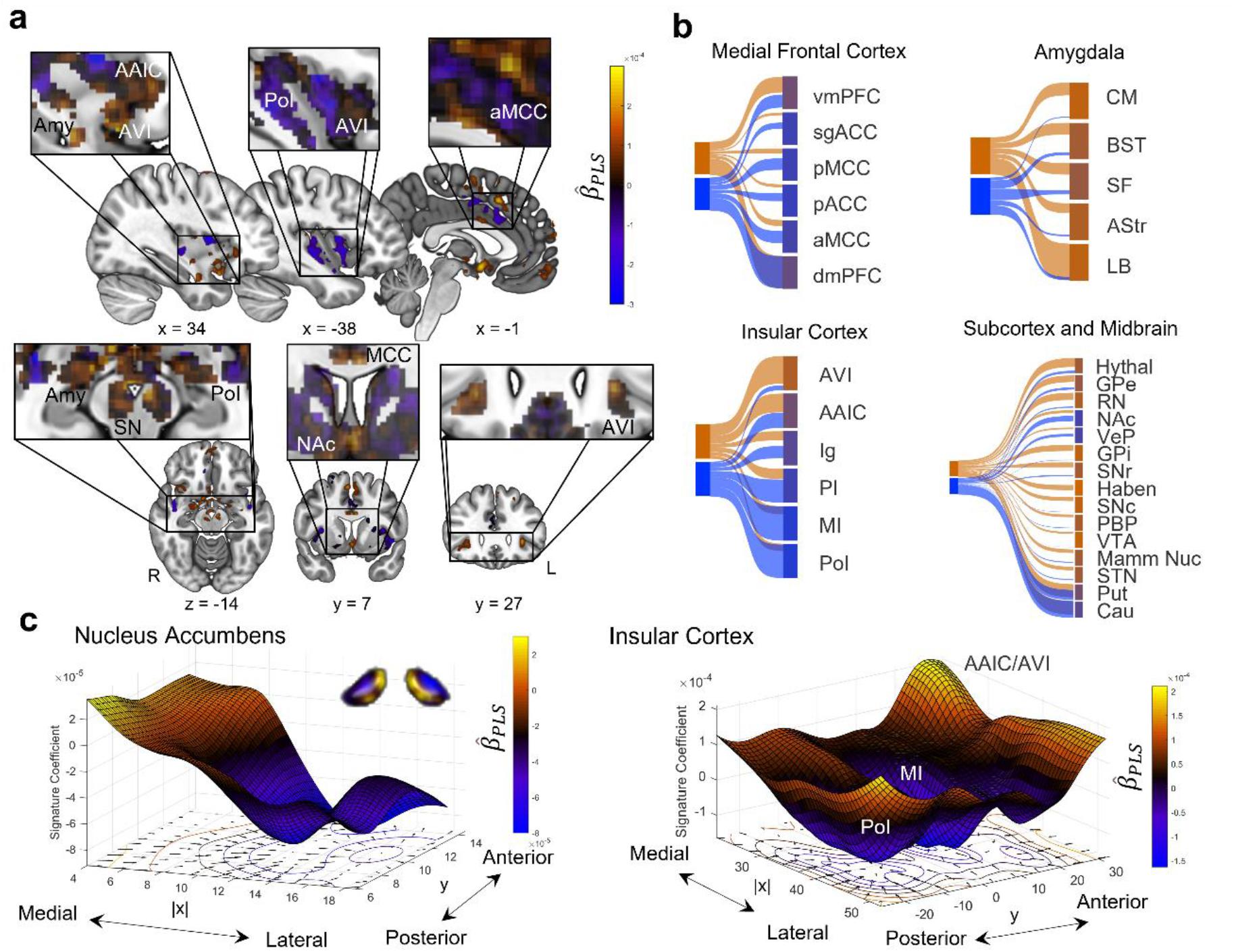
An fMRI-based signature for pleasure. (**a)** Partial Least Squares regression coefficients that define the signature. Warm colors indicate regions in which increases in brain activity contribute to predictions of pleasure, whereas cool colors indicate regions in which brain activity decreases classifications of pleasure. Extreme coefficients (magnitude > .0001) are rendered in volumetric space, and heatmap overlays show unthresholded regression coefficients. (**b**) Alluvial flow plots depict the similarity of coefficients and anatomically defined regions of interest. Positive coefficients are depicted in orange and negative coefficients in blue. **c**, Spatial topography of coefficients in the nucleus accumbens (left) and insular cortex (right). Surfaces depict topographies estimated with fitting thin-plate smoothing splines using the x- and y-coordinates of signature coefficients. Contours and vector fields of the surface gradient in x and y dimensions are depicted below each surface.

Given evidence of both positive and negative signature coefficients in most regions, we next examined whether coefficients exhibited gradients similar to those identified in nonhuman animals. A mediolateral gradient was present in the nucleus accumbens with larger coefficients in more medial portions that decreased laterally (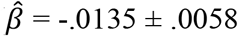 *sem, z* = -2.34, *p* = .0195, linear regression between coefficients and their distance from midline; Fig. 1c). This gradient is consistent with human neuroimaging evidence that rewards activate medial portions of the nucleus accumbens whereas aversive stimuli activate more lateral areas^81,82^. Fine-grained topography was also present within the midcingulate cortex. Coefficients exhibited a clear peak near the bank of the callosal sulcus (*z* = 4.06, MNI_x,y,z_ = [6, 8, 26], *p* < .0001, FDR *q* < .05), whereas predominantly negative coefficients were present throughout the remainder of the midcingulate gyrus. A posterior to anterior gradient was present in the insular cortex (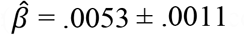 *sem, z* = 4.63, *p* < .0001), with a peak in ventral anterior insula (*z* = 4.80, MNI_x,y,z_ = [-30, 26, -2], *p* < .0001, FDR *q* < .05; Fig. 1c). Rostro-caudal gradients similar to those observed in rodent studies ^11^ were not apparent in the ventral pallidum (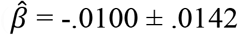 *sem, z* = -.704, *p* = .4815), although the sign and organization of coefficients were roughly consistent with this layout (Supplementary Figure 1). This is possibly due to the small size of this region, spanning roughly 6 mm along its rostro-caudal axis relative to the smoothness of model coefficients, estimated at ∼4.5 mm full-width half-maximum (Supplementary Figure 2)^83^. These findings demonstrate that although the nucleus accumbens, midcingulate, and anterior insula are consistently activated by aversive, rewarding, and salient stimuli^59^, states of pleasure are characterized by unique patterns of activity within each of these regions, consistent with preclinical findings^11,21^.

**Figure 2.**
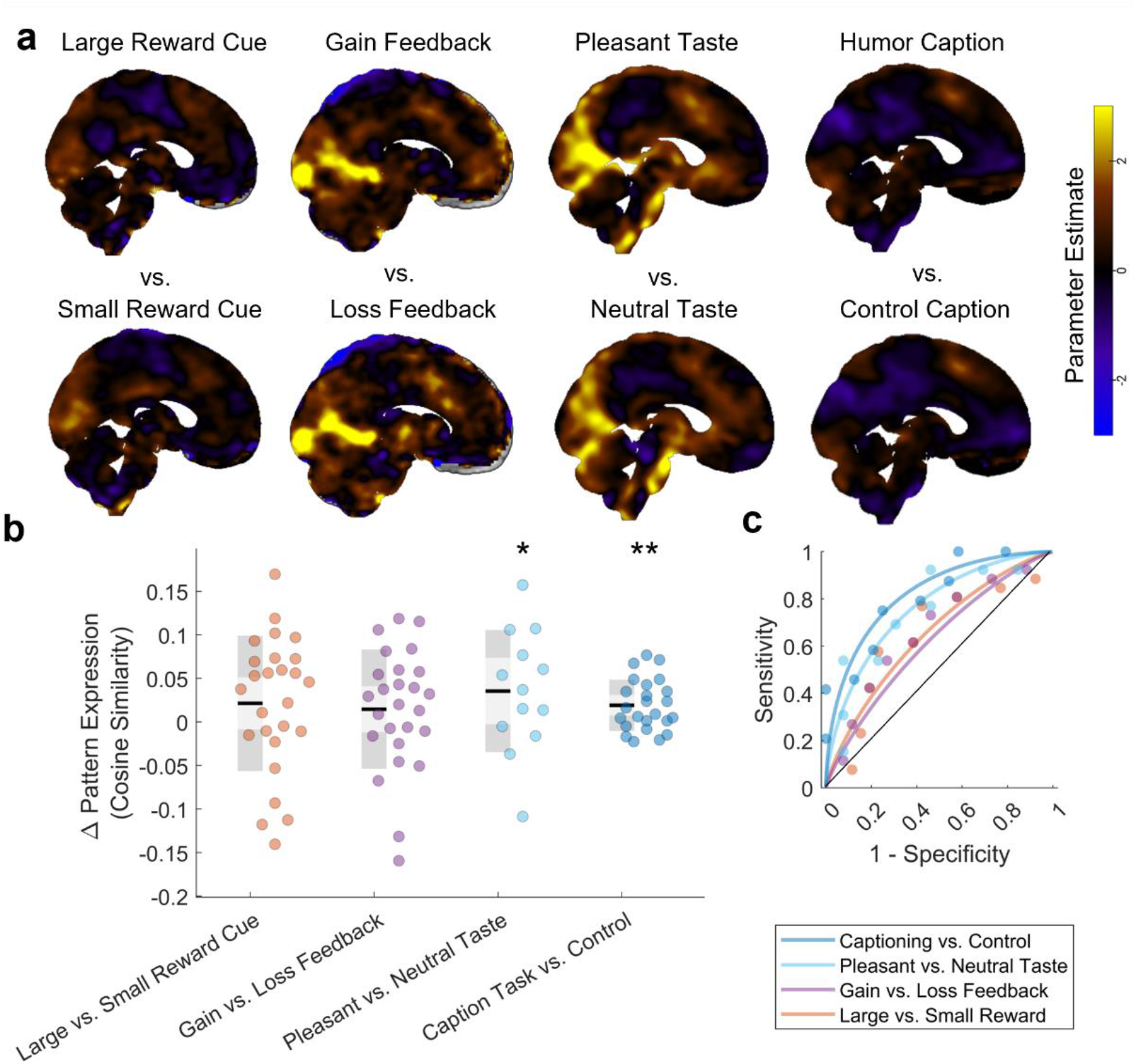
Validation of the pleasure signature. (**a)** Group-average activation maps for four independent validation datasets. Warm colors indicate increases and cool colors indicate decreases in brain activity during each condition compared to baseline. Pairs of experimental conditions are ordered by the predicted difference in signature response. (**b**) Box and whisper plot shows differences in the signature response for each study. Black lines depict the mean response, light shared regions depict one standard deviation, and darker shaded regions two standard errors. Each point corresponds to the response of a single subject. (**c**) Receiver operating characteristic curves for each of the four studies. \**p* < .1, \*\**p* < .05

### Validating the signature in independent studies

Accounts of hedonic function propose that certain stimuli, situations, and behaviors are rewarding because they evoke pleasure^5,6,84^, which is thought to be mediated by a distributed mesocorticolimbic network^11,85^. Evolutionary theories of pleasure go further to suggest that there is a final common pathway in which diverse pleasures are expressed and represented similarly^86^. If the signature we identified captures such a common representation, then it should generalize across instances of pleasure whether they are produced by basic sensory stimuli or more complex cognitive processes. To test this prediction, we evaluated the signature response in multiple independent archival datasets. We first applied the signature to brain activity measured as participants consumed commercial beverages that ranged from hedonically neutral to mildly pleasant^87^. The signature response to the most pleasing beverage, (self-report = 5.32 ± .186 *sem*, pattern response = .0327 ± .0113 *sem*) and a beverage rated as neutral (self-report = 3.93 ± .274 *sem*, pattern response = -0.00265 ± .0146) were discriminable with a medium effect size (AUROC = 0.80, *d* = .72, *p* = 0.0506, Fig. 2), providing initial evidence of generalizability.

In a second generalization test, we examined the signature response during a positive mood induction that combines humor, decision-making, and social feedback^37^. In this task, participants viewed cartoons from the New Yorker Caption Contest and were instructed to select which caption was selected as the winner^88^. Regardless of their selection, participants received feedback that they had chosen the correct option in the majority of trials. We compared the signature response to humor captioning and a matched control task (descriptive captioning) that equated the perceptual, decision-making, and motor aspects of captioning but did not involve humor. The signature response to humor captioning (.137 ± .0119 *sem*) and the control task (.1180 ± .0111 *sem*) were discriminable from one another with a large effect size (AUROC = .82, *d* = 0.92, *p* = .0050), providing further evidence of generalizability.

The sensitivity of the signature to both pleasant tastes and humor suggests it may reflect common coding of pleasure. However, its response to these stimuli could be driven by variables correlated with hedonic impact in the training data, rather than pleasure *per se*. In particular, the signature may have capitalized on signals related to incentive salience and/or reward value ^89,90^ to classify brain states associated with pleasure. To examine this possibility, we tested whether the signature is sensitive to differences in brain activity as participants viewed reward-predictive cues and during reward receipt (i.e., feedback about monetary gains and losses) that differed in terms of reward value but produced only minimal differences in subjective pleasure.

We evaluated the signature response as participants performed an effort-based decision-making task that used visual cues to indicate the magnitude of rewards and the physical effort required to obtain them. We compared responses to cues on low-effort trials (< 30% of maximal effort during a calibration procedure) that indicated participants could receive a large reward ($5) to those indicative of small rewards ($1). Focusing on these conditions ensured that differences in the signature response were related to reward magnitude, rather than the difficulty of the decision or negative value associated with effort. The signature response to large (.0429 ± .0110 *sem*) and small reward cues (.0215 ± .0127 *sem*) were only modestly discriminable from one another, with a small effect size (AUROC = .68, *d* = .39, *p* = .0822).

To further evaluate the possibility that the signature is sensitive to differences in value, we examined its response during a classic reinforcement learning task. In this task, participants learned which of several cues were associated with monetary gains and losses to maximize monetary reward^91^. Computational models of reinforcement learning have demonstrated that momentary changes in positive affect are associated with positive reward prediction errors^92^, and are generally uncorrelated with objective reward magnitude. Based on this evidence, we predicted that if the brain signature responds to the hedonic impact of a stimulus, rather than its value or motivational salience, then it should weakly differentiate visual feedback about gains and losses. Consistent with this prediction, we found the signature response to gains (.0435 ± .0152 *sem, d* = .763) and losses (.0288 ± .0129 *sem, d* = .331) exhibited low levels of discriminability (AUROC = .68, *d* = .30, *p* = .0783).

Across the four validation studies, the signature responded most strongly during states of pleasure (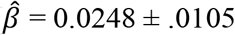 *sem, t*_*173*_ = 2.358, *p* = .0195; linear mixed-effects regression, see Methods). Because functional gradients are a defining feature of hedonic systems, and multivariate predictive models are capable of using information at multiple spatial scales, we additionally tested whether fine-grained patterns within regions that define the signature are necessary for accurate prediction. We constructed a model that replaced each coefficient in the signature with the average of all voxels in each region. Repeating the four validation tests revealed this constrained model did not respond more strongly to states of pleasure than other conditions during manipulations of reward (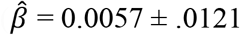 *sem, t*_*173*_ = 0.4681, *p* = .6403, Supplementary Figure 3), demonstrating that variation in fMRI response within regions is necessary for identifying states of pleasure.

To further verify that the pleasure signature is functionally dissociable from evaluative and anticipatory components of reward, we compared the pleasure signature to a recently developed brain signature designed to discriminate between monetary rewards and losses^68^. Whereas the pleasure signature was most sensitive to pleasant taste and humor, the reward signature robustly discriminated gain and loss outcomes during reinforcement learning (AUROC = .85, *d* = .89, *p* = 0.0010) and failed to differentiate states of pleasure from other conditions (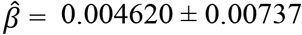 *sem, t*_*173*_ = 0.6271, *p* = 0.5314, Supplemental Fig. 4).

### Cortical and subcortical representation of pleasure

Although it is widely accepted that many sensory cortical areas exhibit a high degree of modularity, i.e., functional specificity^93,94^) that can be consistently detected with fMRI, the extent to which brain areas encoding affect exhibit a similar modular organization is debated^23^. Given that our training and validation studies included a range of reward modalities from primary taste/olfaction to more abstract monetary and humor rewards, we next sought to evaluate the extent to which cortical and subcortical areas contained representations of pleasure that generalized across stimulus types. First, we performed a representational similarity analysis^95^ within the training dataset in regions hypothesized to contain hedonic modules to determine which areas showed modality-specific vs domain-general coding of pleasure. As predicted by the rodent literature^19,20,22^ we observed the clearest evidence for generalizable pleasure coding in the nucleus accumbens (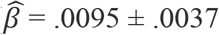 *sem*, z = 2.62, *p* = .0088), ventral pallidum (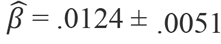 *sem*, z = 2.42, *p* = .0154), and ventromedial prefrontal cortex (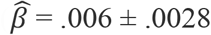 *sem*, z = 2.12, *p* = .0338), with evidence for modality-specific representational geometry within insula and ventromedial prefrontal cortex (Fig. 3).

**Figure 3.**
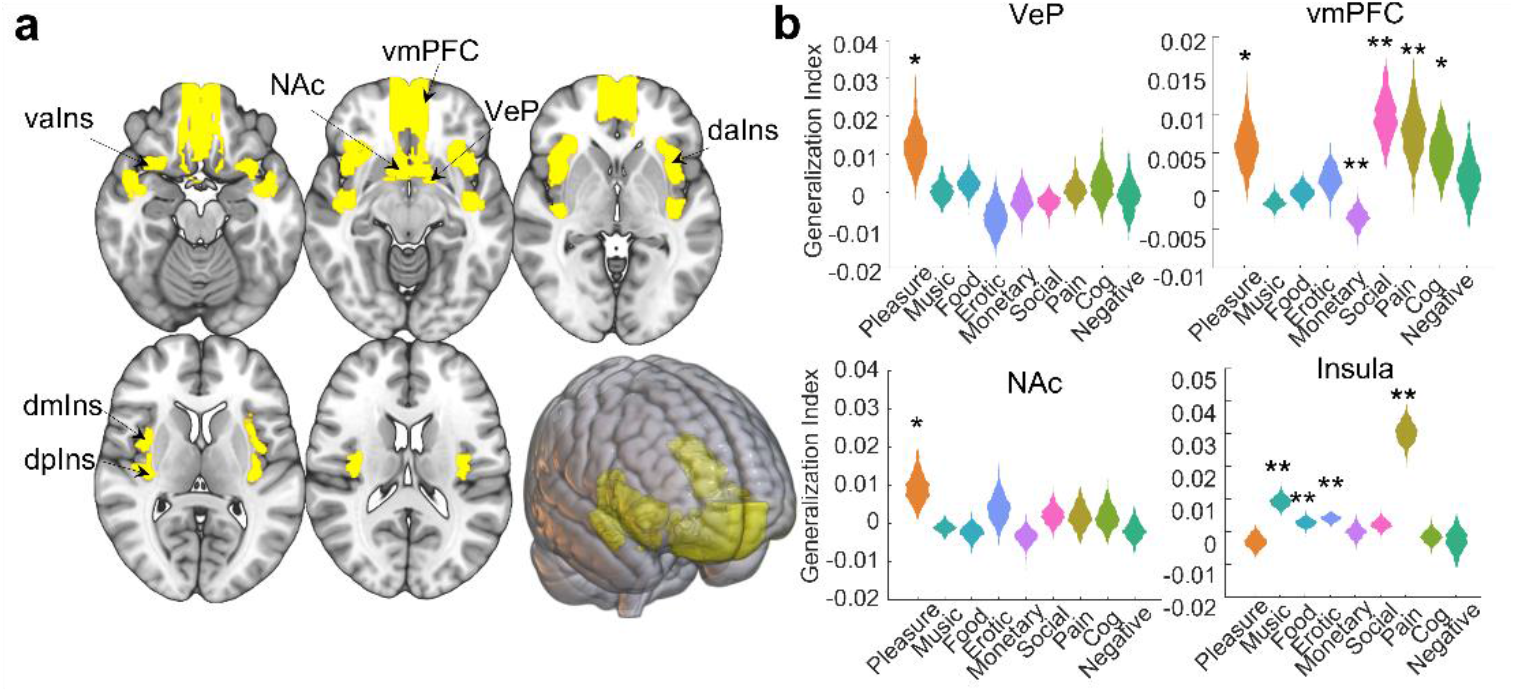
Cortical and subcortical representations of pleasure. (**a**) Volumetric rendering of regions of interest known to contain hedonic hotspots (yellow) on the ICBM152 template. (**b**) Hierarchical representational similarity analysis reveals generalizable patterning related to pleasure in ventral pallidum (VeP), nucleus accumbens (NAc), and ventromedial prefrontal cortex (vmPFC). Representations specific to social pleasure were identified in vmPFC, and representations specific to pleasure induced by music, images of appetizing food, and erotic images were identified in the insula. Bootstrap distributions are shown for the four general domains (pleasure, pain, cognitive control, and negative affect) and five pleasure subdomains (music, food, erotic, monetary, and social pleasure). vaIns = ventral anterior insula, daIns = dorsal anterior insula, dmIns = dorsomedial insula, dpIns = dorsal posterior insula. \**p* < .05, \*\**q*_*FDR*_ < .05

**Figure 3.**
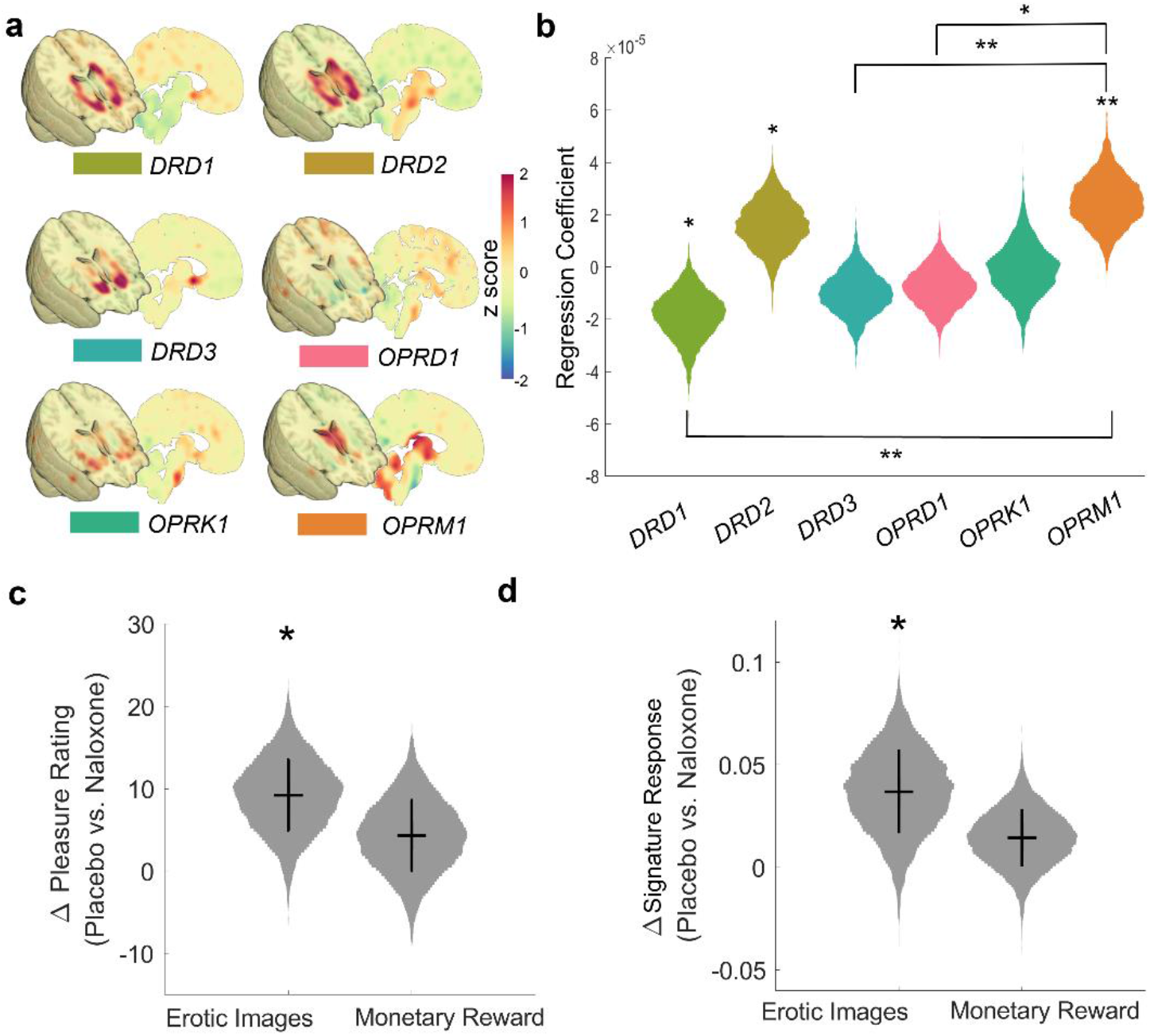
Opioid contributions to the pleasure signature. (**a**) Normalized gene expression maps from the Allen Brain Atlas used to evaluate the spatial correspondence between neurotransmitter gene expression and the pleasure signature coefficients. (**b**) Beta estimates from spatial regression indicate that *DRD2* and *OPRM1* were uniquely associated with coefficients of the pleasure signature. (**c**) Pharmacological challenge using naloxone reduced self-reported pleasure to erotic images, but not monetary rewards. (**d**) Signature response (cosine similarity) was lower with naloxone compared to placebo. Error bars depict the standard error of the mean. Gray shaded regions depict bootstrap distributions (*b* = 10,000). \**p* < .05, \*\**p* < .001

Next, we used our validation datasets to determine the extent to which synthetic “lesions” that excluded signals in cortical and subcortical areas impacted classification of reward-related brain activity. Here, we found that constraining predictive models to exclusively use signals from NAc, VeP, insula, and ventromedial prefrontal cortex had little impact on the discrimination of pleasant taste (ΔAUROC = -.0680, *p* = 0.6501) and substantially impaired humor classification (ΔAUROC = -.3325, *p* = .0010). Conversely, excluding signals from putative hotspot regions did not impair discriminability for pleasant tastes (ΔAUROC = -.0947, *p* = .5515) and resulted in a modest improvement in humor classification (ΔAUROC = 0.1450, *p* = .0470).

These results suggest that—as in rodents—putative hotspots in human NAc, VeP, and OFC show generalizable coding of pleasure across a range of primary and secondary rewards. Indeed, models trained to detect states of pleasure generalized to predict pleasant tastes. However, distributed patterns in these regions were not sufficient to classify humor, which appeared to depend more strongly on the inclusion of medial prefrontal cortex—potentially reflecting the contribution of self-referential processing^10^ or more broadly theory of mind^96^. Taken together, this implies a distributed architecture for pleasure encoding rather than a highly modular organization and highlights distinctions between cortical and subcortical areas for pleasure associated with primary and secondary rewards.

### Evaluating opioid contributions to the signature

Opioidergic mechanisms in mesocorticolimbic structures are thought to play a central role in driving and regulating appetitive behavior, with mu-opioids being particularly involved in hedonic components of reward^11,16-22^. If this is the case, and the signature captures the activity of opioidergic neural populations, there should be a correspondence between the magnitude of signature coefficients and the density of mu-opioid receptors. We tested this hypothesis by performing a spatial regression between the signature coefficients and neurotransmitter gene expression data from the Allen Human Brain Atlas^97^. Due to the considerable overlap of dopaminergic and opioidergic populations in striatum^48,98^, amygdala^99^, and midbrain^100,101^, we included patterns of gene expression for dopamine receptors (*DRD1, DRD2, DRD3*) and opioid receptors (*OPRD1, OPRK1, OPRM1*) in a multiple regression to predict the signature coefficients. Consistent with our hypothesis, this analysis revealed a positive relationship between the spatial profile signature coefficients and *OPRM1* (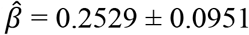 *sem, z* = 2.659, *p* = .0078). Follow-up comparisons revealed this relationship was greater than associations with the expression of other genes, on average (*z* = 2.802, *p* = .0051) although not larger than *DRD2* (*z* = 0.798, *p* = 0.425), Fig. 4, see Supplementary Table 3 for details).

The spatial correspondence between signature coefficients and gene expression is consistent with evidence that opioids mediate human pleasure^102-106^. To assess whether opioids influence the signature response, we tested it on a placebo-controlled cross-over study^107^ using the opioid antagonist naloxone (*n* = 19). In this fMRI study, participants performed an incentive delay task that required participants to make speeded button presses to either obtain monetary rewards or view erotic images. The task was performed in two scanning sessions, concurrent with fMRI and the administration of intravenous naloxone or saline placebo (brain maps showing the effect of naloxone in each condition are shown in Supplementary Figure 5).

Compared to placebo, naloxone reduced self-reported pleasure from erotic images (−9.223 ± 4.436 *sem, z* = -2.130, *p* = .0329, *d* = -.4770), but not monetary rewards (−4.342 ± 4.394 *sem, z* = -.989, *p* = .3229, *d* = -.2267). Naloxone had similar effects on the pleasure signature, reducing its response to erotic images (−0.0376 ± .0191 *sem, z* = -2.037, *p* = .0416, *d* = -.4927), but not to monetary rewards (−0.0168 ± 0.0127 *sem, z* = -1.350, *p* = 0.1771, *d* = -.3215), suggesting that opioids similarly regulate the signature response and pleasure experience, particularly in response to primary rewards.

## Discussion

Diverse forms of human pleasure are thought to be driven by brain systems that originally developed to support the attainment of basic rewards essential for survival—food, social interaction, sex, and maternal care. Under such modular, preadaptation accounts^86^, the same hedonic circuitry that mediates pleasure evoked by basic rewards has been co-opted for more abstract sources of pleasure, such as music, aesthetics, and humor, which additionally involve cortical processing and are heavily influenced by learning. Our findings are broadly consistent with such accounts, as the signature we developed is sensitive to both basic sensory and abstract pleasures, and it does not respond robustly to salient, positive events that lack hedonic impact. Although this suggests the signature may capture activity from a common pleasure pathway, we found that the prediction of humor depended on activity in prefrontal cortex and insula, demonstrating that subcortical modules are insufficient to characterize affective experience in humans.

Following a rich history of attempts to identify neural sources of pleasure^11,108-110^, our label of ‘pleasure signature’ is situated in the context of current theories of hedonic function and neuroimaging data available for training. Eight of the ten studies used to develop the signature verified that participants experienced pleasure using self-reported valence (5 studies) or ratings of the appetitive nature of stimuli (3 studies). The remaining training data comprised brain responses to reward cues (reflecting the magnitude and probability of gains and losses in a mixed gambles task, and the magnitude of immediate and delayed monetary rewards in a delay discounting task) that typically evoke positive affect^111-113^. And even though the training data largely involved visual images (8/10 studies) and passive reward acquisition (8/10 studies), the signature generalized to pleasant taste/olfaction and a novel decision-making task involving humor. As such, the signature we developed is a step towards more precisely characterizing hedonic brain function in humans, taking the critical step of including types of pleasurable stimuli that cannot be easily studied in animals, such as music and humor.

Importantly, the ability of the model to generalize across different types of pleasurable experiences in independent samples does not necessarily suggest that pleasure is undifferentiated or that it is modular in nature. Pleasure is inherently multidimensional, with a hierarchical structure^75^ in which unitary pleasure can be differentiated in terms of antecedent events, sensations, and emotional responses. By design, we trained our model to characterize the apex of this hierarchy so that it would capture generalizable aspects of pleasure rather than stimulus- or situation-specific features. Future work focused on variation within and between different types of pleasure (e.g., sensory, physical, aesthetic, social) is needed to determine how sensory information is transformed into a common representation.

Our results are consistent with neurobiological accounts that characterize affect as an emergent feature of coordinated population activity in distributed neural networks^11^. Rather than requiring only a single region, or depending on large-scale, global signals^69,114,115^, we found that the signature’s ability to accurately predict pleasure was driven by local topography within regions. Although prior meta-analytic summaries have proven invaluable for identifying neural correlates of affect^24-29,59^, these findings suggest that coordinate-based methods lack the precision necessary to discriminate positively and negatively valenced states^59,116^. It will likely be necessary to move from coordinate-based assessments of the literature to pattern-based frameworks to accurately assess the brain basis of affective phenomena.

The spatial layout of the pleasure signature is consistent with prior neuroimaging summaries of positive affect, assessments of mu-opioid receptor availability^117-119^, and observations of hedonic hotspots identified in rodent studies^18,21^. Beyond supporting existing descriptions of pleasure systems, it provides new insight into cortical areas not typically associated with hedonic function. For instance, we observed a peak in signature coefficients in ventral midcingulate cortex (along areas 24a’ and 33’), adjacent to the corpus callosum. Compared to dorsal aspects of the midcingulate, ventral portions of midcingulate have distinct cytoarchitecture^120,121^, functional connectivity^122^, and are not consistently engaged by aversive and cognitively demanding tasks^74,123,124^. Consistent with our findings, stimulation of the callosum near this area has been found to increase spontaneous expression of positive affect in awake humans^125^. Similar to adjacent populations in midcingulate cortex involved in pain affect and reward, hedonic signals could be used to compute the expected value and drive learning about rewards^123,126^, although this possibility remains to be tested.

A somewhat surprising result was our signature’s relatively weak categorization of monetary reward (Fig. 2b). Monetary reward tasks are widely deployed in population neuroimaging research^127-129^ and robustly modulate behavioral responses and corticostriatal activity^130,131^. Many such tasks have been explicitly designed to distinguish neural activity related to reward anticipation or decision-making and activity related to rewarding outcomes (e.g., see refs^132-134^). Importantly, the latter “consummatory” phase of these tasks is often presumed to reflect activity driven by affective responses to monetary reward receipt. Contrary to this interpretation, we found little evidence to suggest that either the anticipatory or consummatory phases of the validation tasks assessed were associated with strong pleasure signals as compared to either primary sensory rewards or secondary pleasure derived from humor. Moreover, an alternative signature developed using a monetary reward task was significantly better at classifying monetary reward from loss (Supplemental Fig. 4). This suggests that monetary reward tasks may primarily capture neural encoding related to reinforcement and instrumental actions, rather than pleasurable affective states. If true, this could have important implications for neuroimaging studies of psychiatric disorders associated with apathy and anhedonia^135,136^.

In sum, the current work identifies a distributed pattern of brain activity that is both sensitive and specific to pleasurable experiences. Strikingly, this signature shares many features with hedonic systems identified in non-human animals, including its anatomical distribution, sensitivity to mu-opioid receptor expression and function, and distinction from non-pleasure reward signals previously linked to dopaminergic pathways. This signature offers new insight into the distributed neural architecture underlying pleasure and can serve as the foundation for more sophisticated models and measures of hedonic function in humans.

## Methods

### Study selection and contrast specification

Because theories of hedonic function focus on the variety of sensory states that can produce similarly pleasurable subjective experiences, we used a meta-analytic approach to develop a generalizable signature for pleasure. This approach enables comparisons between a larger number of experimental conditions and generalization across scanners and populations. To identify a distinct pattern of fMRI activity associated with pleasure, we systematically sampled neuroimaging data from 28 independent studies that manipulated affective valence or engaged cognitive control in healthy individuals (total *n* = 494).

Studies involving manipulations of positive affect were chosen to include five types of reward (images of appetizing food, visual cues of monetary rewards, pleasant music, images and videos depicting pleasant social interactions, and erotic images). Two studies of each reward type were selected for training (10 studies in total, *n* = 224). Activation maps included contrasts between images of appetizing food and baseline^137,138^, the average response to reward cues during a temporal discounting task and baseline^139^, linear variation in reward magnitude during a mixed gambles task^140^, contrasts between pleasant music and resting baseline^141,142^, activation as mothers viewed videos of their infants versus baseline^143^, images of social activities versus baseline following 24 hours of social deprivation^144^, and contrasts between erotic images and baseline^145,146^.

We additionally included activation maps from an archival dataset^74^ of 18 studies (*n* = 270) involving pain, cognitive control, and negative affect. Activation maps from pain studies included contrasts between high (painful) and low (not painful) levels of thermal stimulation^147^, high levels of painful thermal stimulation and baseline^148^, rectal distension trials and baseline^149,150^, and pressure applied to the thumb and baseline^151^. Activation maps from studies involving cognitive control included contrasts between blocks of an *N*-back task and a fixation baseline^152,153^, trials in stop signal tasks compared to baseline^154,155^, and congruent and incongruent trials from studies using the Eriksen Flanker^156^ and Simon^157^ tasks. Activation maps for the negative emotions domain include contrasts between negative and neutral IAPS pictures^158^, negative pictures and baseline^159^, pictures of ex-partners and pictures of close friends^160^, images of others in pain and baseline^161^, and listening to unpleasant affective sounds and baseline^74^.

All participants in the studies listed above provided informed consent in line with local ethics and institutional review boards. Supplementary Table 4 contains descriptions of ethics approvals, image acquisition and analysis, and demographics for each study. Information about data acquisition and experimental paradigms is available in full detail in the corresponding references.

### Feature selection

When selecting features to use for model development, we included regions known to contain hedonic hotspots, namely nucleus accumbens^77^, ventral pallidum^77^, insular cortex^162^, and ventromedial prefrontal cortex including orbitofrontal cortex^74^. We also incorporated several other regions that have connectivity with hedonic hotspots, and several that are involved in processing variables that are often correlated with hedonic impact (e.g., reward value, motivation, attention) and other effortful and/or aversive affective states. These regions include medial prefrontal cortex (subgenual cingulate, perigenual cingulate, anterior midcingulate, posterior midcungulate, ventromedial prefrontal cortex, and dorsomedial prefrontal cortex)^74^, amygdala (central amygdala, basolateral amygdala, superficial amygdala, and amygdalostriatal area)^163^, basal forebrain structures (extendeded amygdala, mammillary nucleus)^77^, striatum (caudate, putamen, globus pallidus)^77^, hypothalamus^77^, habenula^77^, and multiple midbrain nuclei (red nucleus, ventral tegmental area, and substantia nigra)^77^.

### Partial Least Squares specification and estimation

To identify a single pattern of brain activity associated with diverse instances of pleasure that does not respond during other manipulations of affect, we specified a partial least squares (PLS) regression model to predict different levels of a functional hierarchy. The hierarchy included four levels: subject, study, subdomain, and domain. The input data matrix consisted of contrasts from all 494 subjects in the development sample and the output matrix consisted of 46 dummy coded variables (28 studies, 14 subdomains, and 4 domains, with values of +1/–1 based on inclusion/exclusion for each term). PLS models were fit using SIMPLS^164^. A block bootstrap procedure was used for inference on PLS regression coefficients (3,000 samples). In this procedure, observations from individual studies were resampled with replacement to account for dependencies within studies. Normal approximations were made based on the mean and standard deviation bootstrap distributions for each voxel, producing z-maps and associated p-values. PLS regression coefficient maps were thresholded using false discovery rate correction on p-values (two sided) from the bootstrap procedure (*q <* .05).

### Generalization tests

To evaluate the generalizability of the pleasure signature, we examined its response in four independent datasets (Studies 1-4 in Supplementary Table 5), making classifications on the basis of cosine similarity between PLS parameter coefficients estimated during training and test data. The first two of these studies were selected to test model specificity, as they included contrasts that primarily differed in terms of reward value rather than hedonic impact. They included gain versus loss trials in a standard reinforcement learning task^91^ and high versus low reward trials in a task requiring participants to make decisions about expending effort for rewards of varying magnitude^165^. The second pair of studies was chosen to evaluate the sensitivity of the model, as they included contrasts between decision-making about humorous versus non-humorous content^88^ and between pleasant versus neutral tastes^87^. Cohen’s *d* and area under the receiver operating characteristic curve (AUROC) were used to index discrimination within individual studies.

To evaluate the signature response across all four validation studies, a linear mixed effects model was specified with study and a condition × study interaction as fixed effects, and random intercepts for subjects nested within studies. The interaction term in this model was specified to test whether the signature response was larger between conditions for studies that included a manipulation of pleasure (pleasant vs. neutral tastes and humor vs. neutral captioning) and those that did not (large vs. small reward cues and gain vs. loss feedback). The model was fit using maximum likelihood estimation through MATLAB’s fitglme function. Inference was made using a two-sided t-test against zero and confirmed using a parametric bootstrap (10,000 iterations).

### Spatial mappings with neurotransmitter receptor gene expression

We compared the pleasure signature to gene expression maps for dopamine receptors (*DRD1, DRD2, DRD3*) and opioid receptors (*ORPM1, ORPK1*, and *ORPD1*) from the Allen Brain Atlas^97^ accessed in Nifti format through NeuroVault^166^. We performed a multiple regression using the 6 normalized gene expression maps (35,291 voxels by 6 genes) to predict the PLS regression coefficients that define the pleasure signature (35,291 voxels), producing beta estimates that reflect the unique association between gene expression for each receptor type and the pleasure signature. To estimate the variability of betas estimates, a bootstrap procedure was performed using the same bootstrap distribution used to make inference on PLS regression coefficients. Inference was performed using a two-sided test with normal approximation following visual inspection (full distributions are shown in Fig. 3).

### Signature response to naloxone challenge

To further assess the effects of opioids on the pleasure signature, we evaluated its response in a placebo-controlled crossover study^107^ examining the effect of naloxone on mesocorticolimbic activity and self-reported pleasure. As with the other four generalization tests, we computed the cosine similarity between the signature and maps contrasting the placebo manipulation and for erotic images and visual cues that indicating they would receive money after the scanning session. Because some activation maps varied in signal quality and coverage, images that were extreme outliers based on Mahalanobis distance (3 for erotic image contrasts and 2 for monetary reward contrasts) were excluded from this analysis. Including these participants produced slightly smaller yet qualitatively similar estimates of effect size. We computed differences in average pattern expression and self-reported pleasure between the naloxone and saline sessions for both types of stimuli. Bootstrap resampling with normal approximation was performed for both measures, using two-sided tests for inference.

### Representational Similarity Analysis

We constructed model-based representational dissimilarity matrices (RDMs) reflecting the psychological domains and subdomains involved in each study (following methods developed in Kragel et al. 2018^74^, see Supplementary Figure 6). For each RDM, we calculated dissimilarity using 1-Pearson’s *r* correlation distance between multivoxel patterns of brain activity. First, we modeled each of the 28 studies individually to assess differences in pattern generalizability across studies. Then, we modeled the 14 subdomains (food reward, musical reward, monetary reward, social reward, sexual reward, visceral stimulation, thermal stimulation, mechanical stimulation, response conflict, response selection, working memory, visual negative emotion, social negative emotion, and auditory negative emotion) to assess patterns that generalize across studies but differ across subdomains. Lastly, we modeled each of the 4 psychological domains (positive affect, pain, cognitive control, and negative affect) independently to account for response patterns that generalize across studies and subdomains but differ across the four general domains.

We used binary vectors based on study membership to model the observed brain RDMs as a linear combination of individual studies (28 RDMs), subdomains (14 RDMs), and psychological domains (4 RDMs). We then created a set of vectors from the intersubject dissimilarities of these 46 RDMS and a constant RDM, which were used as regressors in a linear regression model. On-diagonal elements, which have zero dissimilarity, were excluded from all analyses. We used a block bootstrap procedure^74^ to obtain *p* values because the general linear model assumes independent errors while dissimilarity matrices exhibit complex dependencies. Positive regression coefficients thus reflect similarities in brain responses that generalize across a psychological domain and cannot be explained by features unique to any subdomain, study, or individual. Two-sided tests were performed for inference.

## Acknowledgements

This project was supported by grants R01MH126083 and R00MH102355 to MTT. DAP was partially supported by grants P50MH119467 and R37MH068376.

## Supplementary Materials

Supplementary Figures 1-6

Supplementary Tables 1-5

## Author Contributions

Conceptualization, methodology, formal analysis, writing – original draft, writing – review and editing, validation, and visualization, PAK; conceptualization, review and editing, and resources,

MTT; writing – review and editing, and resources, RA; writing – review and editing, and resources DAP; conceptualization, writing – original draft, review and editing, and formal analysis, ECH.

## Data Availability

Data used to train the signature will be made available at https://osf.io/, data used for validation are available upon reasonable request.

## Code Availability

Code for all analyses will be made available upon publication at https://github.com/ecco-laboratory/PMA.

