## Supplementary Materials for "A mesocorticolimbic signature of pleasure in the human brain"

This PDF file includes:

Supplementary Figures 1-6
Supplementary Tables 1-5

### Supplementary Figures

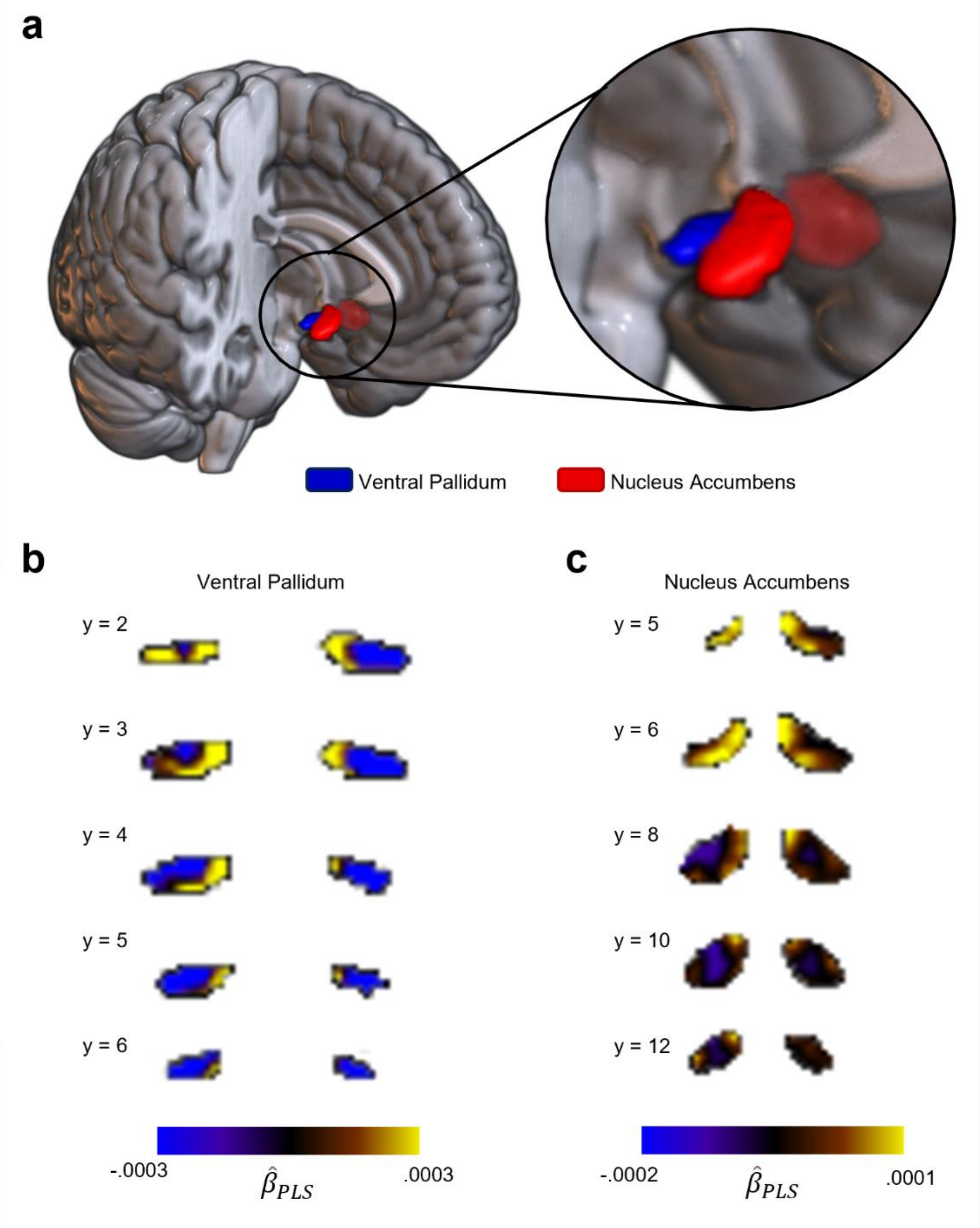

**Supplementary Figure 1.** Signature coefficients within ventral pallidum and nucleus accumbens. (**a**) Volumetric rendering of anatomically defined regions of interest overlaid on the ICBM152 template. (**b**) Signature coefficients (beta estimates from Partial Least Squares regression) within the ventral pallidum that predict states of pleasure. Warm colors are positively associated with predictions of pleasure, whereas cool colors are negatively associated with pleasure. MNI coordinates (mm in the y dimension) are shown next to each section. (**c**) Signature coefficients in the nucleus accumbens.

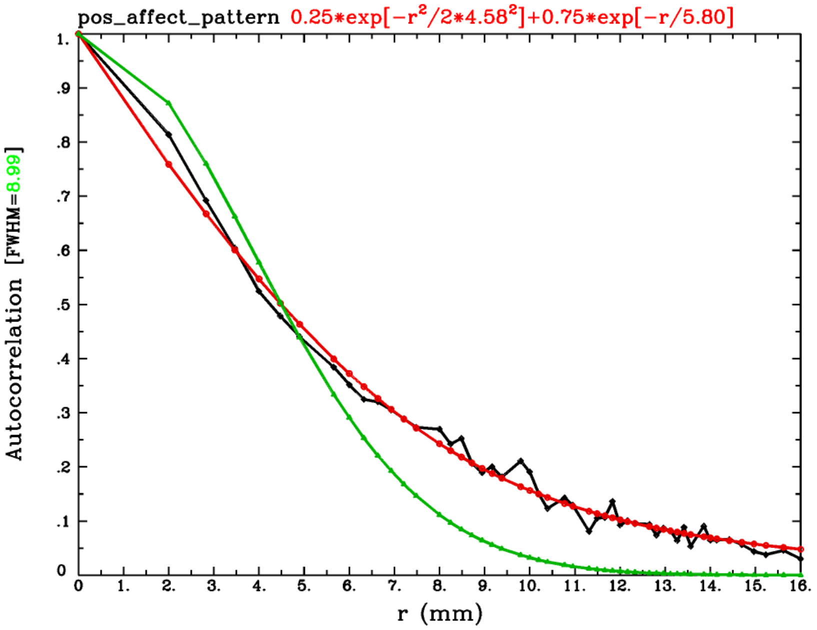

**Supplementary Figure 2.** Estimated spatial smoothness of the pleasure signature. The empirical spatial autocorrelation of the pleasure signature (black) and estimates using both Gaussian (green) and mono-exponential fit (red) are shown. Figure generated from the AFNI program 3dFWHMx.

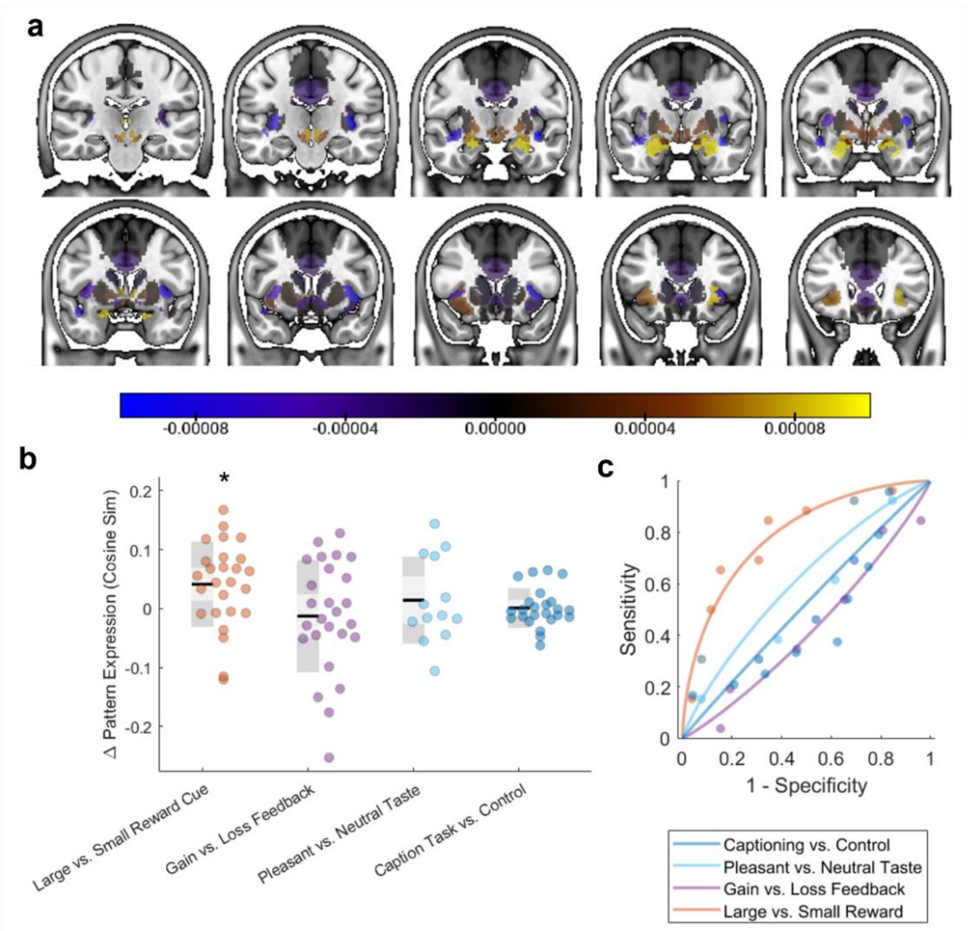

**Supplementary Figure 3.** Simplified region-average model of the signature response is neither sensitive nor specific to pleasure (**a)** Simplified model defined by the average of coefficients from the optimized signature. Warm colors indicate regions in which increases in brain activity contribute to predictions of pleasure, whereas cool colors indicate regions in which increased brain activity leads to fewer classifications of pleasure. (**b**) Box and whisper plot shows differences in the region-average signature response for each study. Black lines depict the mean response, light shared regions depict one standard deviation, and darker shaded regions two standard errors. Each point corresponds to the response of a single subject. (**c**) Receiver operating characteristic curves for each of the four studies. $^*p <$ .01

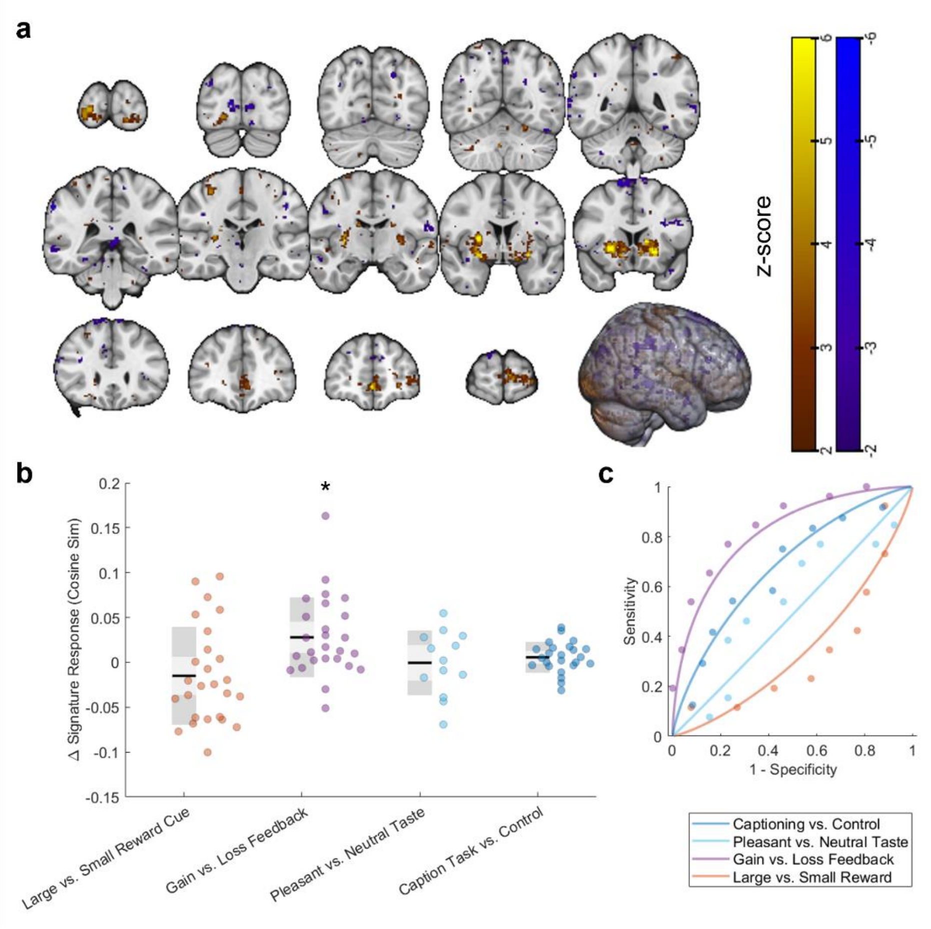

**Supplementary Figure 4.** Brain reward signature is sensitive to reward feedback, but not pleasure. (**a**) Reward signature trained to discriminate gains and losses in a monetary incentive delay task. Warm colors indicate regions in which increases in brain activity contribute to predictions of greater reward, whereas cool colors indicate regions in which increases in brain activity lead to lower levels of reward. (**b)** Box and whisper plot shows differences in the reward signature response for each study. Black lines depict the mean response, light shared regions depict one standard deviation, and darker shaded regions two standard errors. Each point corresponds to the response of a single subject. (**c**) Receiver operating characteristic curves for each of the four studies. *$p < .01$

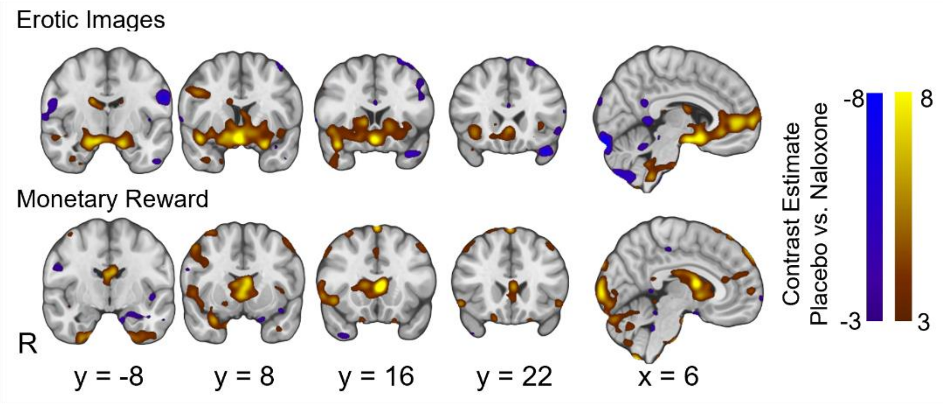

**Figure S5.** Group-average contrast maps showing the effect of naloxone on brain activity during the presentation of erotic images and feedback about monetary rewards. Warm colors indicate greater activity during saline placebo administration compared to naloxone, whereas cool colors indicate a greater response during naloxone administration compared to placebo.

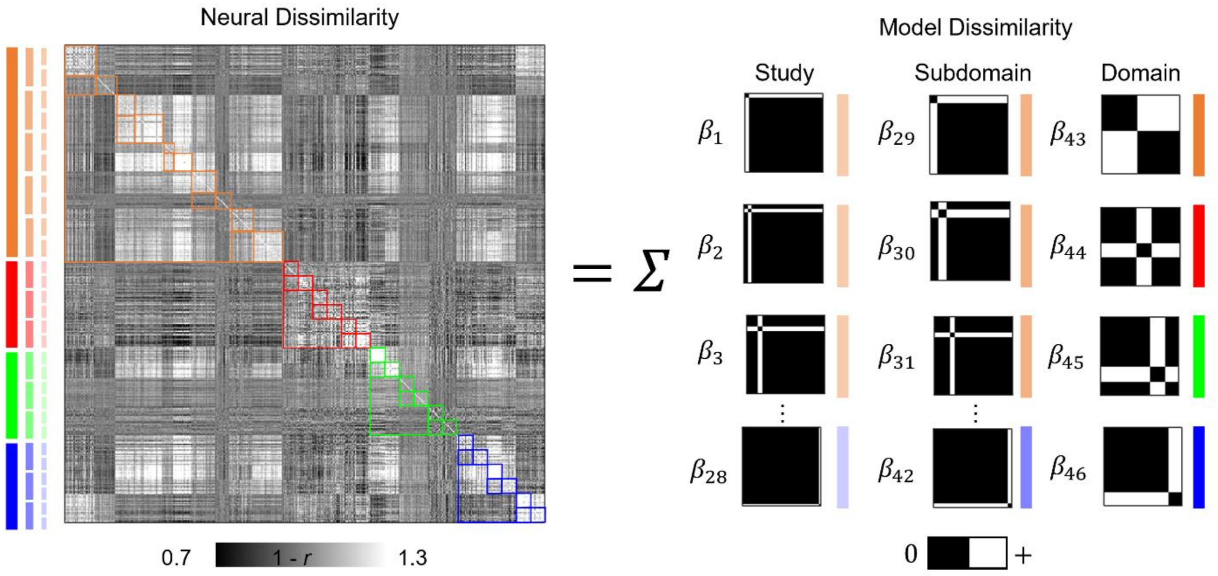

**Supplementary Figure 6.** Decomposing multivariate pattern similarity into study-, subdomain-, and domain-specific components. The matrix in the left panel shows the intersubject dissimilarity of fMRI patterns (n = 494) across all regions of interest. Each element depicts the dissimilarity (1 – Pearson′s correlation coefficient) in brain activity patterns for two individuals. Colored bars to the left indicate corresponding levels in a functional hierarchy including positive affect, pain, cognitive control, and negative affect. The right panel shows how the observed neural dissimilarity across pairs of images from the 28 studies is modeled as a weighted summation of dummy coded dissimilarity matrices specified according to study (28 parameters), subdomain (14 parameters), and domain (4 parameters) membership, in addition to a constant term (not shown). Adapted from Figure 1 of Kragel et al. 2018.

### Supplementary Tables

#### Supplementary Table 1. Peak Signature Coefficients

| *Positive Coefficients* | | | | | |
|---|---|---|---|---|---|
| *Region* | *Volume (cubic mm)* | *$MNI_{x,y,z}$* | | | *Peak Z* |
| Habenula | 192 | -4 | -24 | 0 | 4.894 |
| Anterior Ventral Insula | 552 | -30 | 26 | -2 | 4.8032 |
| Dorsomedial Prefrontal Cortex | 784 | 0 | 16 | 46 | 4.3539 |
| Midcingulate Cortex (Area 33) | 256 | -4 | 6 | 26 | 4.2934 |
| Internal Globus Pallidus | 64 | -14 | -4 | -4 | 4.1182 |
| Red Nucleus | 240 | 0 | -18 | -10 | 4.1022 |
| Midcingulate Cortex (Area 33) | 208 | 6 | 8 | 26 | 4.06 |
| External Globus Pallidus | 64 | -14 | 0 | 0 | 4.0464 |
| Caudate Tail | 64 | 30 | -34 | 6 | 3.8734 |
| Subthalamic Nucleus | 112 | 14 | -12 | -6 | 3.853 |
| Premotor Cortex (Area 24) | 64 | -10 | -22 | 46 | 3.8432 |
| Premotor Cortex (Area 6a) | 96 | -24 | -10 | 46 | 3.7777 |
| Putamen | 64 | 24 | -8 | 12 | 3.7552 |
| Caudate Tail | 64 | 30 | -30 | -4 | 3.7501 |
| Internal Globus Pallidus | 64 | 18 | -10 | -2 | 3.7347 |
| Amygdala | 64 | 24 | -4 | -16 | 3.7245 |
| Dorsomedial Prefrontal Cortex (Area 8) | 64 | -14 | 34 | 52 | 3.7223 |
| Substantia Nigra | 96 | -8 | -18 | -14 | 3.7011 |
| Medial Frontal Cortex | 128 | 14 | 16 | 38 | 3.6996 |
| Posterior Insula | 64 | 34 | -8 | -10 | 3.6862 |
| Basal Forebrain | 64 | -6 | 2 | 4 | 3.6637 |
| Anterior Ventral Insula | 64 | 34 | 28 | -4 | 3.6569 |
| Anterior Cingulate Cortex | 64 | 8 | 14 | 24 | 3.6278 |
| *Negative Coefficients* | | | | | |
| *Region* | *Volume (cubic mm)* | *$MNI_{x,y,z}$* | | | *Peak Z* |
| Posterior Insula | 920 | 40 | -12 | -8 | -4.8465 |
| Posterior Insula | 448 | -44 | 6 | -6 | -4.7464 |
| Premotor Cortex (Area 6ma) | 272 | 26 | 14 | 62 | -4.74 |
| Premotor Cortex (Area 6ma) | 448 | 20 | -6 | 70 | -4.4971 |
| Posterior Insula | 192 | -40 | -16 | -10 | -4.446 |
| Dorsal Caudate | 144 | 12 | -6 | 24 | -4.3029 |
| Middle Insula | 352 | 36 | 6 | 8 | -4.1944 |
| Middle Insula | 376 | -34 | 4 | 8 | -4.052 |
| Premotor Cortex (Area 6ma) | 176 | 20 | 4 | 66 | -4.025 |
| Dorsal Caudate | 96 | 18 | 2 | 24 | -3.9974 |
| Premotor Cortex (Area 6mp) | 160 | -12 | -8 | 72 | -3.9713 |
| Supplementary Motor Area | 96 | 4 | -8 | 64 | -3.8666 |
| Posterior Midcingulate | 248 | 0 | -6 | 42 | -3.8009 |
| Premotor Cortex (Area 6) | 64 | 18 | 24 | 60 | -3.6688 |
| Premotor Cortex (Area 24) | 64 | 4 | -16 | 50 | -3.6416 |
| Premotor Cortex (Area 6mp) | 64 | 2 | -12 | 66 | -3.6355 |
| Posterior Insula | 88 | -38 | -18 | -6 | -3.6123 |

**Supplementary Table 2. Spatial similarity of signature coefficients and regions of interest**

| *Region* | *Mean Pos* | *SEM Pos* | *Z Pos* | *p Pos* | *Mean Neg* | *SEM Neg* | *Z Neg* | *p Neg* | *Mean Diff* | *SEM Diff* | *Z Diff* | *p Diff* |
|---|---|---|---|---|---|---|---|---|---|---|---|---|
| Posterior Midcingulate Cortex | 0.080 | 0.016 | 5.112 | 0.000 | 0.178 | 0.031 | 5.689 | 0.000 | -0.098 | 0.043 | -2.308 | 0.021 |
| Anterior Midcingulate Cortex | 0.099 | 0.018 | 5.500 | 0.000 | 0.177 | 0.038 | 4.714 | 0.000 | -0.078 | 0.052 | -1.505 | 0.132 |
| Perigenual Anterior Cingulate Cortex | 0.054 | 0.017 | 3.273 | 0.001 | 0.128 | 0.024 | 5.256 | 0.000 | -0.074 | 0.038 | -1.955 | 0.051 |
| Subgenual Anterior Cingulate Cortex | 0.051 | 0.013 | 3.854 | 0.000 | 0.081 | 0.022 | 3.689 | 0.000 | -0.030 | 0.031 | -0.971 | 0.332 |
| Ventromedial Prefrontal Cortex | 0.250 | 0.028 | 8.827 | 0.000 | 0.200 | 0.028 | 7.021 | 0.000 | 0.050 | 0.046 | 1.079 | 0.281 |
| Dorsomedial Prefrontal Cortex | 0.490 | 0.030 | 16.525 | 0.000 | 0.483 | 0.030 | 16.200 | 0.000 | 0.007 | 0.049 | 0.146 | 0.884 |
| Right Posterior Insular Area 1 | 0.037 | 0.009 | 3.979 | 0.000 | 0.135 | 0.022 | 5.992 | 0.000 | -0.098 | 0.027 | -3.595 | 0.000 |
| Left Posterior Insular Area 1 | 0.025 | 0.009 | 2.868 | 0.004 | 0.137 | 0.031 | 4.401 | 0.000 | -0.111 | 0.037 | -3.043 | 0.002 |
| Right Para-Insular Area | 0.033 | 0.011 | 3.020 | 0.003 | 0.140 | 0.027 | 5.226 | 0.000 | -0.107 | 0.033 | -3.244 | 0.001 |
| Left Para-Insular Area | 0.056 | 0.017 | 3.248 | 0.001 | 0.055 | 0.013 | 4.382 | 0.000 | 0.001 | 0.026 | 0.023 | 0.981 |
| Right Middle Insular Area | 0.038 | 0.011 | 3.611 | 0.000 | 0.124 | 0.027 | 4.574 | 0.000 | -0.085 | 0.036 | -2.386 | 0.017 |
| Left Middle Insular Area | 0.035 | 0.013 | 2.631 | 0.009 | 0.164 | 0.027 | 6.023 | 0.000 | -0.129 | 0.037 | -3.501 | 0.000 |
| Right Insular Granular Complex | 0.038 | 0.012 | 3.285 | 0.001 | 0.052 | 0.017 | 2.972 | 0.003 | -0.014 | 0.026 | -0.524 | 0.600 |
| Left Insular Granular Complex | 0.027 | 0.010 | 2.718 | 0.007 | 0.069 | 0.016 | 4.200 | 0.000 | -0.042 | 0.023 | -1.772 | 0.076 |
| Right Anterior Ventral Insular Area | 0.112 | 0.024 | 4.669 | 0.000 | 0.028 | 0.012 | 2.397 | 0.017 | 0.084 | 0.032 | 2.615 | 0.009 |
| Left Anterior Ventral Insular Area | 0.121 | 0.024 | 4.979 | 0.000 | 0.018 | 0.008 | 2.280 | 0.023 | 0.103 | 0.029 | 3.547 | 0.000 |
| Right Anterior Agranular Insula Complex | 0.098 | 0.020 | 4.887 | 0.000 | 0.043 | 0.012 | 3.674 | 0.000 | 0.055 | 0.028 | 1.958 | 0.050 |
| Left Anterior Agranular Insula Complex | 0.065 | 0.018 | 3.518 | 0.000 | 0.071 | 0.017 | 4.265 | 0.000 | -0.006 | 0.031 | -0.201 | 0.841 |
| Putamen | 0.130 | 0.038 | 3.455 | 0.001 | 0.132 | 0.034 | 3.823 | 0.000 | -0.002 | 0.069 | -0.023 | 0.982 |
| Caudate Nucleus | 0.153 | 0.027 | 5.641 | 0.000 | 0.172 | 0.031 | 5.585 | 0.000 | -0.019 | 0.053 | -0.361 | 0.718 |
| Ventral Pallidum | 0.021 | 0.007 | 2.905 | 0.004 | 0.032 | 0.011 | 2.782 | 0.005 | -0.010 | 0.014 | -0.729 | 0.466 |
| Internal Globus Pallidus | 0.060 | 0.011 | 5.639 | 0.000 | 0.012 | 0.006 | 1.973 | 0.049 | 0.047 | 0.016 | 3.061 | 0.002 |
| External Globus Pallidus | 0.067 | 0.011 | 5.892 | 0.000 | 0.030 | 0.009 | 3.210 | 0.001 | 0.036 | 0.019 | 1.929 | 0.054 |
| Nucleus Accumbens | 0.041 | 0.013 | 3.196 | 0.001 | 0.053 | 0.017 | 3.210 | 0.001 | -0.012 | 0.026 | -0.463 | 0.643 |
| Hypothalamus | 0.112 | 0.025 | 4.410 | 0.000 | 0.040 | 0.016 | 2.447 | 0.014 | 0.072 | 0.036 | 1.972 | 0.049 |
| Habenula | 0.044 | 0.010 | 4.354 | 0.000 | 0.000 | 0.001 | 0.261 | 0.794 | 0.044 | 0.011 | 4.121 | 0.000 |
| Extended Amygdala | 0.069 | 0.015 | 4.577 | 0.000 | 0.020 | 0.010 | 1.900 | 0.057 | 0.049 | 0.021 | 2.345 | 0.019 |
| Superficial Amygdala | 0.077 | 0.020 | 3.779 | 0.000 | 0.026 | 0.014 | 1.843 | 0.065 | 0.051 | 0.031 | 1.643 | 0.100 |
| Basolateral Amygdala | 0.204 | 0.039 | 5.277 | 0.000 | 0.030 | 0.014 | 2.220 | 0.026 | 0.174 | 0.050 | 3.475 | 0.001 |
| Centromedial Amygdala | 0.075 | 0.022 | 3.377 | 0.001 | 0.016 | 0.011 | 1.415 | 0.157 | 0.059 | 0.031 | 1.916 | 0.055 |
| Amygdalostriatal Area | 0.053 | 0.013 | 4.115 | 0.000 | 0.013 | 0.007 | 1.800 | 0.072 | 0.040 | 0.018 | 2.241 | 0.025 |
| Ventral Tegmental Area | 0.020 | 0.007 | 2.715 | 0.007 | 0.002 | 0.003 | 0.848 | 0.397 | 0.017 | 0.009 | 1.816 | 0.069 |
| Subthalamic Nucleus | 0.039 | 0.009 | 4.412 | 0.000 | 0.011 | 0.006 | 1.953 | 0.051 | 0.028 | 0.013 | 2.201 | 0.028 |
| Substantia Nigra pars reticulata | 0.055 | 0.011 | 4.823 | 0.000 | 0.017 | 0.007 | 2.431 | 0.015 | 0.038 | 0.016 | 2.344 | 0.019 |
| Substantia Nigra pars compacta | 0.040 | 0.009 | 4.297 | 0.000 | 0.007 | 0.004 | 1.614 | 0.106 | 0.033 | 0.012 | 2.711 | 0.007 |
| Red Nucleus | 0.073 | 0.016 | 4.684 | 0.000 | 0.016 | 0.008 | 2.062 | 0.039 | 0.057 | 0.021 | 2.732 | 0.006 |
| Parabrachial Pigmented Nucleus | 0.028 | 0.008 | 3.672 | 0.000 | 0.008 | 0.004 | 2.020 | 0.043 | 0.020 | 0.011 | 1.857 | 0.063 |
| Mammillary Nucleus | 0.043 | 0.019 | 2.270 | 0.023 | 0.014 | 0.010 | 1.335 | 0.182 | 0.030 | 0.026 | 1.144 | 0.253 |

Cosine similarity of pleasure signature and binary masks of regions of interest. The mean, standard error of the mean, z-scores and p-values estimated from bootstrap resampling of subjects within studies (3,000 iterations) are shown.

**Supplementary Table 3. Spatial regression and correlations of signature coefficients and gene expression**

| *Gene* | *Mean* | *Bootstrap SE* | *Z score* | *p-value* |
|---|---|---|---|---|
| *DRD1* | -0.019 | 0.009 | -2.060 | 0.039 |
| *DRD2* | 0.016 | 0.009 | 1.793 | 0.073 |
| *DRD3* | -0.010 | 0.008 | -1.276 | 0.202 |
| *OPRD1* | -0.008 | 0.008 | -1.006 | 0.314 |
| *OPRK1* | -0.001 | 0.010 | -0.085 | 0.932 |
| *OPRM1* | 0.025 | 0.010 | 2.659 | 0.008 |

### Supplementary Table 4. Training datasets

| *Study Number* | *Reference* | *Subdomain* | *Domain* | *Stimulus/ Paradigm* | *Contrast*[†] | *Number of Runs* | *Run Length* | *Total Number of Trials* | *Stimulus Duration* | *N (female)* | *Mean Age* | *IRB/Ethics Approval Committee* | *MRI System* |
|---|---|---|---|---|---|---|---|---|---|---|---|---|---|
| 1 | Iranpour et al. 2015 | Food Reward | Positive Affect | Images of food and nonfood | Appetitive food vs baseline | 2 | 9’ | 108 | 2.5 s | 20 (11) | 25.5 | University Hospital of Clermont-Ferrand | 3T GE Discovery MR750 |
| 2 | Smeets et al. 2013 | Food Reward | Positive Affect | Images of food and nonfood | Appetitive food vs baseline | 1 | 10’ | 128 | 2.5 s | 30 (30) | 22.1 | University Medical Center Utrecht, The Netherlands | 3T Phillips Achieva |
| 3 | Castrellon et al. 2019 | Monetary Reward | Positive Affect | Temporal Discounting | Reward cues vs baseline | 2 | ~8’44” | 84 | ~10 s | 25 (13) | 20.9 | Vanderbilt University | 3T Phillips Intera Achieva |
| 4 | Tom et al. 2007 | Monetary Reward | Positive Affect | Mixed Gambles | Parametric modulation of reward | 3 | 8’ | 256 | 3 s | 16 (9) | 22 | University of California, Los Angeles | 3T Siemens AG Allegra |
| 5 | Kragel et al. 2015 | Music Reward | Positive Affect | Music Clips | Pleasant music vs baseline | 7 | 12’ | 28 | 2.2 m | 32 (19) | 26 | Duke University | 3T GE MR750 |
| 6 | Lepping et al. 2016 | Music Reward | Positive Affect | Music Clips | Pleasant music vs baseline | 5 | ~5’24” | 90 | 33 s | 20 (11) | 28.5 | University of Kansas Medical Center | 3T Siemens Skyra |
| 7 | Laurent et al. 2018 | Social Reward | Positive Affect | Videos of infants | Videos of own baby vs baseline | 2 | 7’30” | 72 | 15 s | 25 (25) | 26.4 | University of Oregon | 3T Siemens Allegra 3 |
| 8 | Tomova et al. 2020 | Social Reward | Positive Affect | Images of food, social situations | Social images vs baseline | 6 | ~4’54” | 108 | 5 s | 40 (27) | 26 | Massachusetts Institute of Technology | 3T Siemens Prisma |
| 9 | Dalenberg et al. 2018 | Sexual Reward | Positive Affect | Erotic Images | Erotic images vs baseline | 1 | 20’ | 40 | 4 s | 21 (0) | 24.39 | University Medical Center Groningen | 3T Phillips Intera |
| 10 | Kragel et al. 2019 | Sexual Reward | Positive Affect | Images from IAPS | Erotic images vs baseline | 2 | 7’30” | 112 | 4 s | 18 (10) | 25 | University of Colorado Boulder | 3T Siemens TrioTIM |
| 11 | Atlas et al. 2010 | Cutaneous Somatic Pain | Pain | Thermal stimulation | High vs low pain | 8 | 6’18” | 64 | 10 s | 19 (9) | 25.5 | Columbia University | 1.5T GE Signa TwinSpeed Excite HD |
| 12 | Wager et al. 2013 | Cutaneous Somatic Pain | Pain | Thermal stimulation | 49.3° C vs baseline | 10 | ~6’40” | 8 | 12.5 s | 33 (22) | 27.9 | Columbia University | 1.5T GE Signa TwinSpeed Excite HD |
| 13 | Kano et al. 2017 | Visceral Pain | Pain | Rectal distention | Distension vs baseline | 6 | 12’ | 36 | 18 s | 29 (15) | 22.5 | Tohoku University | 3T Siemens TrioTIM |
| 14 | Rubio et al. 2015 | Visceral Pain | Pain | Rectal distention | Distension vs baseline | 6 | 12’ | 36 | 18 s | 15 (9) | 24* | Comité de Protection des Personnes Sud Est V, France | 3T Philips Achieva TX |
| 15 | Čeko et al. 2022 | Mechanical Pain | Pain | Pressure Stimulation | 7 kg/cm$^2$ vs baseline | 4 | 6’37” | 8 | 10 s | 15 (4) | 26.9 | University of Colorado Boulder | 3T Siemens TrioTIM |
| 16 | Čeko et al. 2022 | Mechanical Pain | Pain | Pressure Stimulation | 4, 5, 6 and 7 kg/cm$^2$ vs baseline | 6 | 7’3” | 24 | 10 s | 15 (8) | 24.2 | University of Colorado Boulder | 3T Siemens Prisma |
| 17 | DeYoung et al. 2009 | Working Memory | Cognitive Control | N-back (faces and words) | 3-back blocks vs baseline | 6 | 5’52” | 348 | 2 s | 104 (59) | 22.7 | Washington University Medical Center | 3T Siemens Allegra |
| 18 | van Ast et al. 2016 | Working Memory | Cognitive Control | N-back (words) | N-back blocks vs baseline | 4 | ~5’ | 306 | 2 s | 21 (10) | 22.2 | Columbia University | 3T Philips Achieva |
| 19 | Aron et al. 2007 | Response Selection | Cognitive Control | Stop signal Task | All trials vs baseline | 6 | 5’32" | 768 | ~1 s | 15 (5) | 28.1 | UCLA | 3T Siemens Allegra |
| 20 | Xue et al. 2008 | Response Selection | Cognitive Control | Stop signal Task | All trials vs baseline | 6 | 6'4" | 768 | ~1 s | 15 (9) | 23.6 | UCLA | 3T Siemens Allegra |
| 21 | ds101*** | Response Conflict | Cognitive Control | Simon Task | Incongruent vs Congruent Trials | 2 | 5' | 24 | 1 s | 21 (9) | 30.5 | New York University | 3T Siemens Allegra |

| | | | | | | | | | | | | | |
|---|---|---|---|---|---|---|---|---|---|---|---|---|---|
| 22 | Kelly et al. 2008 | Response Conflict | Cognitive Control | Eriksen Flanker Task | Incongruent vs Congruent Trials | 2 | 10' | 12 | 2 s | 26 (10) | 28.3 | New York University | 3T Siemens Allegra |
| 23 | Gianaros et al. 2014 | Visual Emotion | Negative Emotion | Images from IAPS | Negative pictures vs baseline | 1 | 11'28" | 15 | 7 s | 183 (88) | 42.7 | University of Pittsburgh | 3T Siemens TrioTIM |
| 24 | Yarkoni et al. 2011** | Visual Emotion | Negative Emotion | Images from IAPS | Negative vs neutral pictures | NR | NR | NR | ~4 s | 108 (NR) | NR | Stanford University, Columbia University | 1.5 and 3T GE Signa LX Horizon Echospeed, 1.5T GE Signa Twin Speed Excite HD scanner |
| 25 | Kross et al. 2011 | Social Emotion | Negative Emotion | Images of ex-partners | Images of ex-partner vs friend | 2 | ~13’30” | 16 | 15 s | 40 (21) | 20.8 | Columbia University | 1.5T GE Signa TwinSpeed |
| 26 | Krishnan et al. 2016 | Social Emotion | Negative Emotion | Images of others in pain | High pain pictures vs baseline | 9 | 8’ | 27 | 11 s | 30 (12) | 25.2 | University of Colorado Boulder | 3T Siemens TrioTIM |
| 27 | Kragel et al. 2018 | Auditory Emotion | Negative Emotion | Sounds from IADS | Unpleasant Sounds vs baseline | 4 | 10’ | 12 | 8 s | 15 (7) | 31.1 | University of Colorado Boulder | 3T Siemens TrioTIM |
| 28 | Kragel et al. 2018 | Auditory Emotion | Negative Emotion | Sounds from IADS | Unpleasant Sounds vs baseline | 4 | 9’18” | 12 | 8 s | 15 (9) | 24.4 | University of Colorado Boulder | 3T Siemens TrioTIM |

Note. Informed consent was provided for each study listed. Data for studies 11-28 were accessed from Neurovault at https://neurovault.org/collections/3324/. * Median age ** Data are collapsed across four separate studies *** Data not previously published. † Baseline involved eyes open fixation for all studies. NR = Data not reported. IAPS = International Affective Picture System. IADS = International Affective Digital Sounds

**Supplementary Table 5. Validation datasets.**

| *Study Number* | *Reference* | *Stimulus/ Paradigm* | *Contrasts†* | *Number of Runs* | *Run Length* | *Total Number of Trials* | *Stimulus Duration* | *N (female)* | *Mean Age* | *IRB/Ethics Approval Committee* | *MRI System* |
|---|---|---|---|---|---|---|---|---|---|---|---|
| 1 | Dalenberg et al. 2017 | Tasting beverages | Pleasant vs neutral beverages | 3 | ~15’ | 36 | 3.5 s | 45 (0) | 24.05 | University Medical Center Groningen | 3T Philips Intera |
| 2 | Admon et al. 2015 | Descriptive comic captioning | Humorous vs neutral captions | 4 | ~15’ | 72 | 6 s | 30 (22) | 30.2 | McLean Hospital | 3T Siemens TrioTIM |
| 3 | Arulpragasam et al. 2018 | Effort Based Decision Making | High vs low monetary reward | 2 | ~ 9’ | 16 | ~3 s | 28 (15) | 20.2 | Emory University | 3T Siemens TrioTIM |
| 4 | *** | Reinforcement Learning | Monetary gains vs losses | 1 | 10’ | 72 | 3 s | 26 (17) | 27.7 | Emory University | 3T Siemens TrioTIM |
| 5 | Buchel et al. 2018 | Monetary Incentive Delay | Saline vs naloxone | 5 | ~ 12’ | 110 | 1.5 s | 19 (0) | 25.48 | University Medical Center Hamburg-Eppendorf | 3T Siemens TrioTIM |
| 6 | Gordon et al. 2017 | Resting State | NA | 10 | ~ 30’ | NA | NA | 10 (5) | 29.1 | Washington University School of Medicine | 3T Siemens TrioTIM |

Note. Informed consent was provided for each study listed. *** Data not previously published. † Baseline involved eyes open fixation for all studies.
